# Development and Optimisation of in vitro Sonodynamic Therapy for Glioblastoma

**DOI:** 10.1101/2023.06.19.545530

**Authors:** Andrew Keenlyside, Theodore Marples, Zifan Gao, Hong Hu, Lynden Guy Nicely, Joaquina Nogales, Han Li, Lisa Landgraf, Anna Solth, Andreas Melzer, Kismet Hossain-Ibrahim, Zhihong Huang, Sourav Banerjee, James Joseph

## Abstract

**Background:** Sonodynamic therapy (SDT) is currently on critical path for glioblastoma therapeutics. However, the molecular mechanisms underlying the therapeutic functions of SDT remain enigmatic due to the lack of intricately optimised instrumentation capable of modulating SDT delivery in vitro. We established and validated an automated in vitro SDT system and assessed its SDT efficacy.

**Methods:** An automated in vitro SDT system was established and validated to allow the application and mapping of focused ultrasound fields under varied acoustic exposure conditions and setup configurations. Ultrasound field simulations were performed to assess the acoustic energy exposure. Systematic in vitro investigations were performed to optimise ultrasound frequency, intensity, plate base material, thermal effect, and the integration of live cells.

**Results:** In the presence of 5-ALA, focused ultrasound induces apoptotic cell death in primary patient-derived glioma cells with concurrent upregulation of intracellular reactive oxygen species. Intriguingly, primary glioma stem neurospheres also exhibit remarkably reduced 3D growth upon SDT exposure. Further, ultrasound field simulations performed established acoustic field distribution and the impact of acoustic standing waves within the well-plates.

**Conclusions:** The optimised in vitro SDT system and associated ultrasound field simulations establish the basis for further understanding of SDT and its therapeutic potentials in cancer.

## Introduction

Glioblastoma (GBM) is a highly invasive and refractory grade IV glioma with extremely poor median patient survival of 12-15 months from initial diagnosis ^1,2^. A GBM tumour is hypoxic with microenvironment niches exhibiting considerable genetic instability and neovascularisations leading to highly heterogenous tumour progression^3^. The tumour comprises primarily of glioma stem cells which exhibit a diverse transcriptome with remarkable reciprocity which grants effective resistance to many traditional and novel treatment options ^4,5^. Complete surgical resection is often impossible as the tumour invades healthy brain by hijacking normal neuronal functions ^6^. However, surgical interventions have improved over the past years, especially with the FDA approval of 5-Aminolevulinic Acid (5-ALA) in 2017, a photosensitizer that allows surgeons to delineate normal brain tissue from glioblastoma.

5-ALA is a metabolite in the heme pathway which is upregulated in GBM ^7^. Upon oral intake, exogenous 5-ALA crosses the inflamed blood-brain barrier (BBB) and selectively accumulates in the tumour ^8^. GBM-specific upregulation of the heme pathway promotes rapid metabolic breakdown of 5-ALA into the photosensitiser protoporphyrin IX (PpIX) ^9^ which exhibits a selective accumulation within the tumour ^10^. PpIX fluorescence (405-633nm^9^), can provide effective intra-operative guidance for tumour resection and progression-free survival ^11,12^. However, tumours eventually recur, predominantly within 2mm of even maximally resected tumour margins^13^. Recent studies have suggested exploiting the selective accumulation of photosensitisers to induce death in residual tumour cells using photodynamic therapy (PDT)^14^. However, PDT requires a craniotomy for effective tumour targeting using laser light to activate 5-ALA, thus ruling out regular repeated dosing. Interestingly, in 2014, 5-ALA was reported to be a sonosensitiser which opened up the possibility of non-invasive excitation of the 5-ALA-metabolite PpIX using focused ultrasound ^15 16^. Focused ultrasound has been FDA and EMA-approved and in use for hyperthermic thalamotomy for essential tremor and BBB modulation ^17,18^. When applied together with sonosensitisers, this is referred to as sonodynamic therapy (SDT).

The use of SDT as a cancer therapy procedure is under investigation. However, the mechanism of FUS-mediated photoexcitation of PpIX remains unknown. Several mechanisms may contribute to activation ^16^. Mechanisms involving microbubble collapse generating blue channel photoemissions and direct mechanical effects have been discussed^19^. These photoemissions cause the development of primarily mitochondrial singlet reactive oxygen species (ROS)^20^. Subsequent cell cycle arrest and caspase activation cause the selective death of tumour cells, sparing normal neural cells which do not contain elevated PpIX concentrations ^21^. Multiple other cytotoxic pathways may be implicated, with different GBM subclones showing variable responses to ROS^22,23^.

Recently, SDT studies showed positive outcomes from in vivo and clinical settings. Significant improvements to survival and tumour inhibition were seen in SDT-treated glioma rat models utilising a 20-minute sonication (1.06MHz, 5.5W/cm^2^) with +2.5°C of sub-ablative tumour hyperthermia^24^. Rat models observed greater benefit when using pulsed FUS treatments (86ms, 8.6% duty cycle) ^25^. These successes led to the instigation of the first phase-0 human trials (clinical trial ID: NCT04559685). However, the exact molecular mechanisms leading to this combined effect of FUS and sonosensitisers in malignant glioma remains unclear.

No optimum in vitro setup and related cell sampling techniques for in vitro SDT has been systematically optimised. Additionally, the effect of SDT on primary patient-derived glioma stem cells in vitro has never been studied. This paper optimises various systems parameters such as the plate material and thickness, acoustic intensity, thermal effects, ultrasound frequency, and methods to reduce pressure distortion due to the formation of acoustic standing waves and eventually reports systematic cell-based studies to validate the experimental SDT system. Taken together, the current study reports a comprehensive system for in vitro SDT and establishes the first systematic testing and cell death investigations in state-of-the-art primary patient derived glioma cell and stem cell models.

## Methods and Materials

### The SDT experimental setup

Figure 1 illustrates the experimental setup. The experimental SDT system comprised of an open-top acrylic water bath (423mm x 486mm x 302mm) and an external water loop with an in-line water heater (T08200 ETH 300W, Hydor) and pump (LET 775, 12V, 65PSI, Lei Te Co.) to circulate temperature-controlled water. The design featured two motorised scanning platforms, one with three degrees-of-freedom, and another with two degrees. The former scanning platform was used to perform US field mapping across X, Y and Z axes using a needle hydrophone (D=1mm, Precision Acoustics), mounted onto a custom 3D printed bracket. Hydrophone output signals were acquired using a high-resolution oscilloscope (PicoScope® 4224, Pico Technology) and relayed for data logging using a custom MATLAB programme. Well plates (*Standard clear 96-well plate with 1mm polystyrene-base*, ThermoFisher Scientific, product ID: 167008 or a ND *Ultrasound-translucent 96-well plate:* (μClear®, 190μm film-based), Greiner Bio-One, product ID: 655097) were mounted using custom, 3D printed, brackets to the latter stage and had motorised range in the X and Y axes. This bracket could be manually adjusted to position the wells in the focal range of the ultrasound transducer. Both stages were translated using motorised linear slides (Motorized XSlide™, XN10 Series, Lead Screw, Velmex Inc.) driven by stepper motor controllers (VXM™-2 Motor Controller, Velmex Inc.) actioned by a custom MATLAB programme. A thermal camera (thermoIMAGER TIM 160, Micro-Epsilon UK Ltd.) was mounted co-axially with the transducer and manually focused (supplementary Fig S4A). Two ultrasound transducers of differing frequencies (666kHz, Precision Acoustics UK, and 1193kHz, homemade) were used. These were interchangeable prior to testing and were mounted in the water bath underneath the stages. The ultrasound transducers were driven using a signal generator (AFG3101, Single Channel, Arbitrary/Function Generator, Tektronix UK Ltd.).

**Figure 1.**
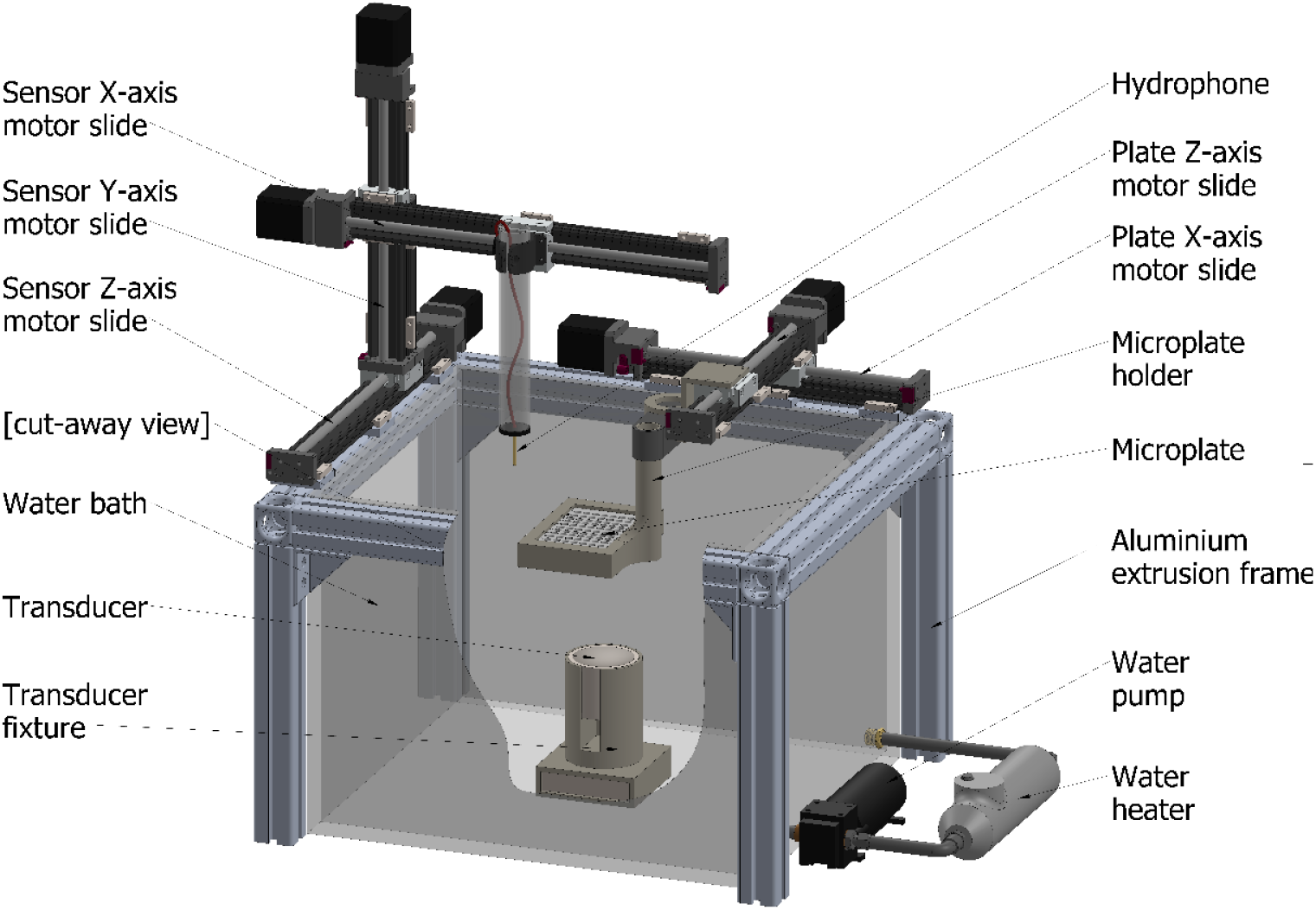
Illustration of the in vitro experimental setup: water bath, needle hydrophone, plate holder and 96-well plate, transducer and fixture, sensor motor slides for the X, Y, and Z axis, and water heater and pump.

### Needle Hydrophone field mapping

A needle hydrophone was used to map a 2D area of the ultrasound field to sample the distribution of acoustic intensity within the system. The hydrophone recorded the pressure between incremental movements (0.1mm) (X-axis) in one direction, before elevating by one increment and repeating this process in the opposite direction. This continued until the desired data grid on an XY plane was formed, with an equal resolution of height and width. This process was executed using a custom programme written in MATLAB. Heatmap of acoustic intensity across the target areas were developed to visualise pressure distribution This protocol was applied to the following experiments, except for *In well (PVA – Submerged)*, to create field mapping in all three cardinal planes. Four distinct exposure setups were devised to examine the effects of altered system parameters:

#### No well plate

The well plate was removed from the testing setup to allow the open field to be mapped within the water bath, with the water level well above the focal point of the US field. The focal point was identified manually and also served as a reference point for the centre of the mapped area (supplementary Fig. S1A)

#### In well (water level)

The well plate and thermal camera were included in the testing setup and the stage was adjusted to place the well plate at the level of the water surface and with the focal point of the ultrasound within the target well. The thermal camera was focussed on the target well which was filled with 400μL water. Surrounding wells contained 100μL of phosphate buffered solution (PBS) (supplementary Fig. S1B).

#### In well (submerged)

The well plate was included in the testing setup and the stage was adjusted to place the well plate 60mm below the water surface and with the focal point of the ultrasound within the target well (supplementary Fig. S1C).

#### In well (PVA – Submerged)

PVA negative casts of the wells were made to study the effects of different ultrasound reflection coefficients within the wells. The PVA protocol and setup is descried in the supplementary methods.

### 2D Cell culture reagents and assays

Primary patient-derived glioblastoma cell GBM22 was acquired from the Mayo Clinic, USA and cultured as reported previously^26,27^. 5-Aminolevulinic acid hydrochloride >98% (5-ALA) (Sigma Aldrich, St. Louis, Missouri, United States) was used for cell dosing alongside standard GBM media. GBM22 cells were plated with 50uL of standard GBM media in standard clear tissue-culture treated 96-well plates described above or 18-well chambered coverslips (Ibidi, Munich, Germany). 5-ALA was diluted in GBM media to 2mM and added to cells at a final concentration of 1mM 2 hours prior to sonication. Non-ALA treated controls were supplemented with 50μL of GBM media alone. For the assessment of reactive oxygen species, CellROX Green (ThermoFisher Scientific, Waltham, MA, United States) was added to both 5-ALA stock solutions and GBM media intended for non-ALA conditions as per manufacturer’s instructions. To assess apoptosis, Annexin V FITC (Abcam, Cambridge, UK) was used. Using suction, all media was removed from the wells within 15 minutes of the completion of sonication. The wells were washed with 50μL buffer solution, which was then removed before the addition of 25μL of buffer solution containing Annexin V-FITC diluted as par manufacturer’s instructions. The cells were incubated for 90 minutes prior to microscopy. Annexin V and CellROX agents were never used together. Bright field and fluorescent imaging were taken using a Thermo Scientific EVOS imaging system.

### 3D glioma stem neurosphere culture and treatments

GBM120 3D glioma stem neurosphere line were acquired from the Mayo Clinic, USA and cultured as reported previously ^27^. Neurosphere cultures, for either formation or formed assay, were plated with poly-D-lysine (PDL) (ThermoFisher Scientific, Waltham, Massachusetts, United States) to ensure adherence to the base of the well of the 96-well plate. Cultures were then dosed using the above 5-ALA method and treated for 30s cumulative sonication at standard parameters. No fluorescent reagents/dyes were used in these cultures. Bright field images of the neurospheres were taken at day 0 (pre-treatment) and day 1 onwards every 3-4 days until 21 days using a Thermo Scientific EVOS imaging system. Formation assays were treated 1 day after plating whilst formed assays were treated after 21 days of culture development and monitored for a further 21 days.

### Sonication of cells

Sonication of cell lines followed a standardised set of US exposure parameters: 0.4w/cm ^2^, 10% duty cycle, 90ms pulse length. These were continued until either 30s or 60s of cumulative sonication had been completed (5 or 10 minutes of treatment respectively) unless otherwise stated in the figure legends. The water bath was maintained at 37°C. These tests were conducted in TC treated polystyrene plates using the *In well (water level)* setup and 666kHz transducer. The overall energy applied to cells at the centre of each well was estimated to be 1.5J (for 30s cumulative sonication).

### Antibodies and immunoblotting

1.5hr post treatment, the cells were lysed in 1x western blot loading buffer (5x: Tris pH 6.8; SDS 10%; glycerol 30%; bromophenol blue 0.02%; beta-mercaptoethanol 5%). Buffer components were purchased from Sigma Millipore. Immunoblotting was carried out as discussed previously ^28 29^ using the following antibodies: total AKT (Cell Signaling #9272), phospho Thr308 AKT (Cell Signaling #2965), phospho Ser473 AKT (Cell Signaling #9018), phospho Thr202/Tyr204 p44/42 ERK1/2 (Cell Signaling #4376), and GAPDH (Cell Signaling #5174).

### System calibration

The SDT system was calibrated to ensure accurate FUS dose estimation and concordance to pressure field simulations. The system can be calibrated manually or automatically to accelerate testing protocols with negligible difference when the transducer position remained constant (Supplementary Fig. S2). The diameter of the focus point may be altered by transducer frequency and curvature of the transducer surface. When a 666kHz transducer is utilised, the focus point diameter was measured to be 4.5mm, slightly narrower than the inner diameter of a typical 96-well plate well. The effect on adjacent wells is minimal (Supplementary Figs. S3).

### Simulation and Data Analysis

US field simulation models based on finite element analysis were developed in COMSOL Multiphysics v5.4 (ANSYS Inc.). The setup components were modelled using their realistic properties sourced from the data sheets of each product. The model was meshed to a maximum element size of 0.6mm for each simulation. The “pressure acoustics, frequency domain” model was used to generate the acoustic physics of the transducer. Analysis of hydrophone field mapping data and the production of relevant figures was completed in MATLAB (MathWorks, Massachusetts, USA). Image J (National Institute of Health, Maryland, USA) was used to assess neurosphere diameter. PRISM – GraphPad (California, USA) was used to generate data characteristics and apply statistical analyses.

## Results

### The use of pressure field simulations

Pressure field simulations were initially performed to optimise the experimental setups and estimate the US exposure levels on the cells and assess the impact of materials in an in vitro setup. Field simulations were also used where field mapping using a hydrophone was not feasible or would alter the pressure field. For field simulations to be a useful tool they must be concordant with the hydrophone field mapping approach to provide comparable results. Simulations and field mapping of three different test setups were undertaken to measure the concordance of the results (Figure 2). The setups tested were, 1) transducer field without a plate or any materials in place, 2) in-well testing of a submerged plate 60mm below the water bath surface, and 3) in-well testing of a plate resting on the water level with 400μL of water in each well.

**Figure 2.**
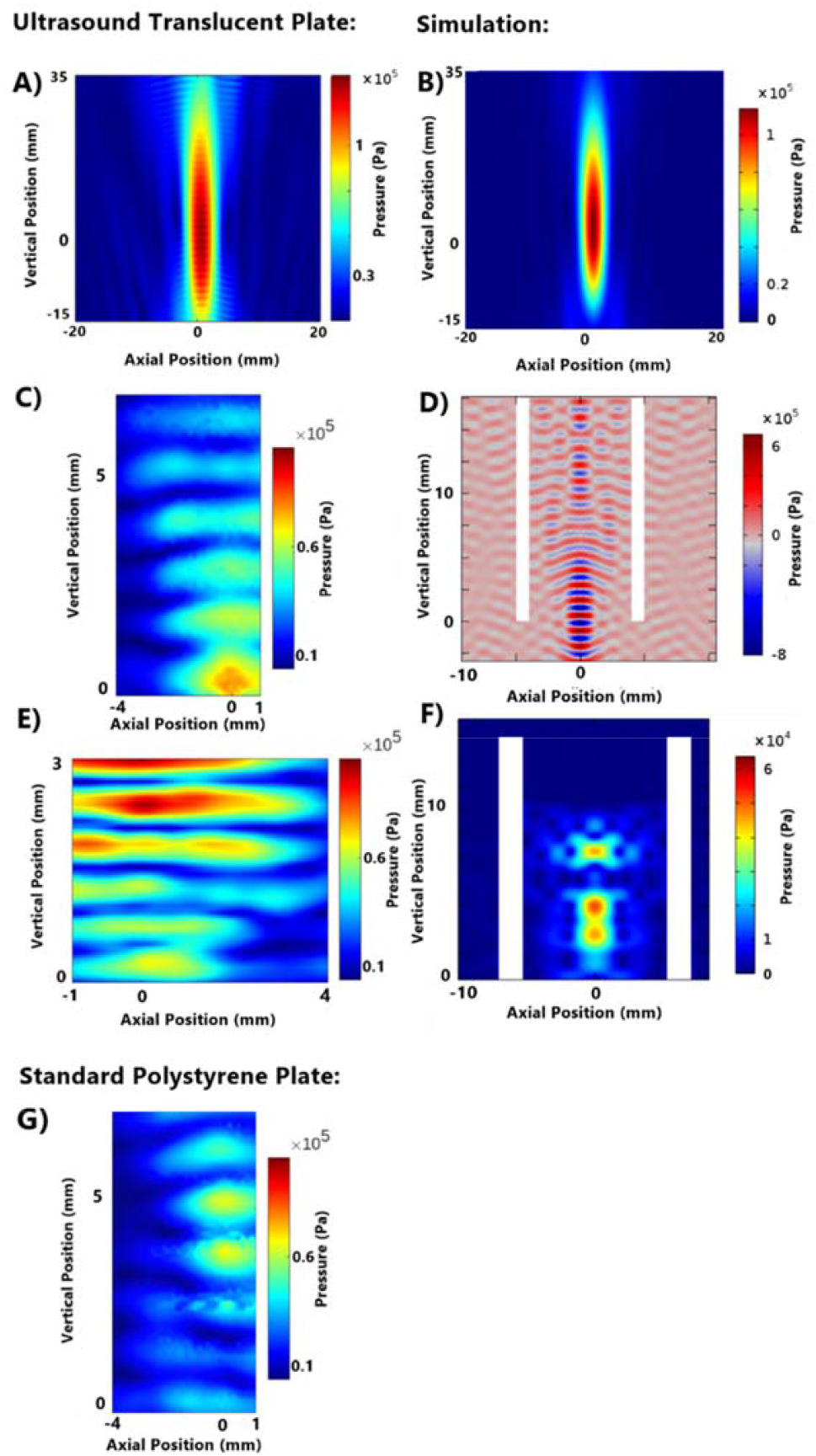
Comparison between ultrasound translucent plate hydrophone mapping and simulation for no well plate, in-well (submerged), and in-well (water level) setups and standard polystyrene plate in-well (submerged setup). 0.4w/cm ^2^ continuous wave. Simulations completed in COMSOL. A) Hydrophone field mapping: no well plate B) Field Simulation: no well plate C) Hydrophone field mapping: in-well (submerged) in an ultrasound translucent plate (190μm film-base). D) Field Simulation: in-well (submerged) E) Hydrophone field mapping: in-well (water level) F) Field simulation: in-well (water level). G) Hydrophone field mapping: in-well (submerged) in a standard polystyrene plate (1mm base thickness).

Simulation results showed strong concordance to in vitro field mapping (Fig 2). In-well (submerged) and in-well (water level) hydrophone field maps gave comparatively limited field mapping capabilities within the wells, when compared to simulation, due to the risk of contact damage to the needle hydrophone. In both hydrophone mapping and simulation, in-well (submerged) study showed a peak pressure at the base of the well which then weakens. Simulations also showed an increase in pressure levels further above the level visible in hydrophone mapping, highlighting the expanded map that simulations provide. In-well (water level) simulations assumed media filled adjacent wells and showed consistent properties of pressure dampening across the lateral walls of the well plate. This can be contrasted against hydrophone field mapping which illustrates the effect, until 1mm above the base of the well. This change is due to the fluid level in the adjacent well containing 100uL compared to the 400uL in the target well required for hydrophone mapping.

### Plate Base Material and Thickness

Standard tissue culture treated sterile plates made from clear moulded polystyrene were used. The base of the well had an average thickness of 1mm. Acoustic boundaries formed due to the finite thickness of the well base resulted in the distortion of the pressure field within the well. Therefore, well plates with thinner bases were considered as a potential alternative for performing in vitro focused ultrasound studies. The in-well (submerged) ultrasound fields of both standard polystyrene 96-well plates and ultrasound translucent (microclear) plates, were compared to determine the impact of base polystyrene on ultrasound field distortion within a well. Continuous wave FUS at 5.5w/cm ^2^ was used for the studies. Hydrophone pressure field mappings performed on both plates under the described conditions showed clear differences in US field distribution within the standard and microclear well plates (Fig 2).

Investigations were also performed to assess the field distribution in the region immediately above the base of the well, where the cell cultures are fixed. Increased acoustic translucency was observed in the microclear plates (Fig 3) due to its thinner base. Horizontal axis profiles showed reduced variance in acoustic intensity across the base of standard plates compared to microclear plates (23.7% and 59.2% respectively). However, microclear plates showed lower variance in the vertical axis or when the radius of the horizontal axis was reduced to 0.8mm, to resemble the US beam, compared to standard plates (38.1% vs. 42.9%; 18.9%)

**Figure 3.**
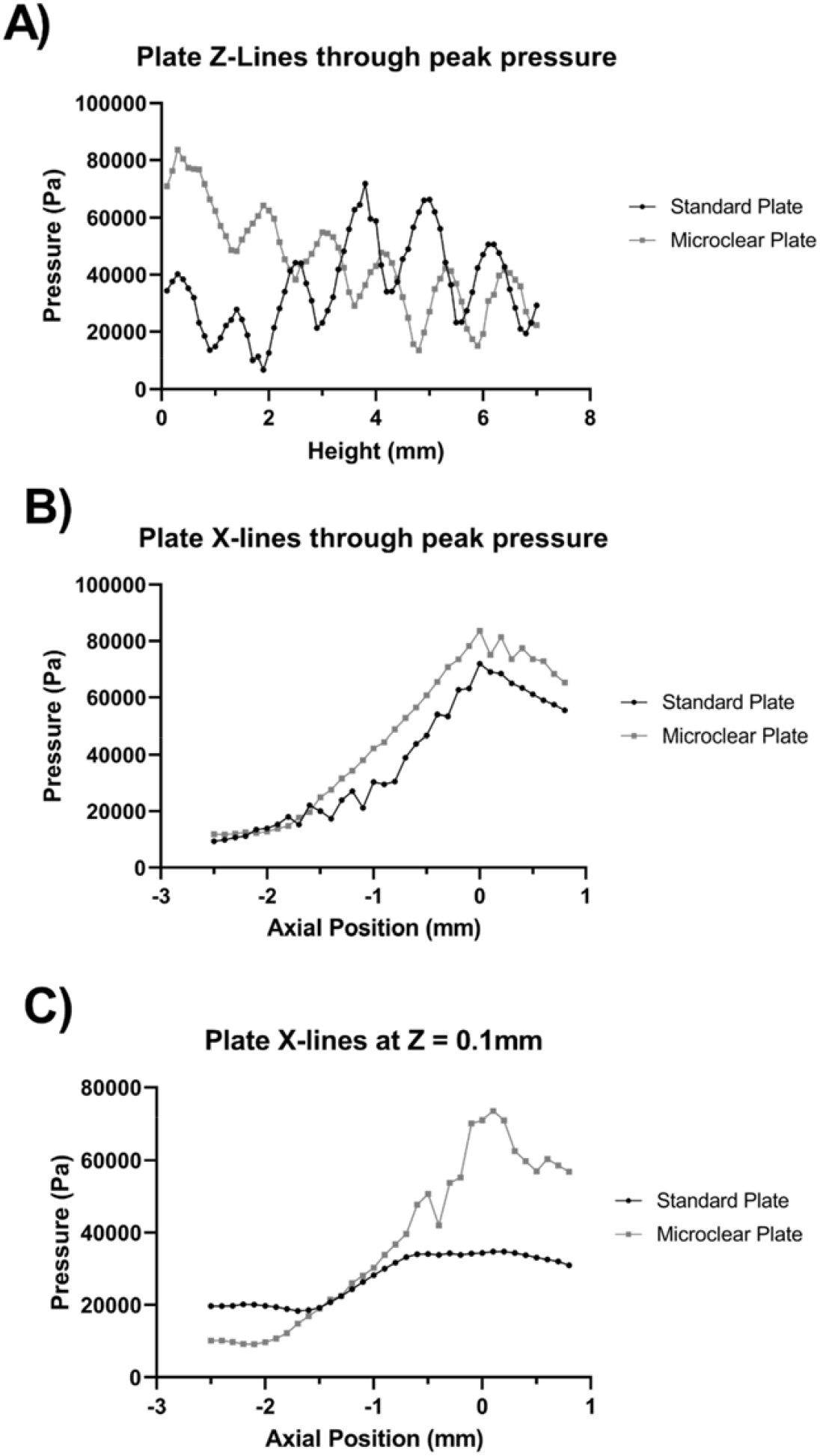
Single axis pressure profiles to compare standard polystyrene and ultrasound translucent (MicroClear) plates. A) Z-axis profiles intercepting the peak pressure point. B) X-axis profiles intercepting the peak pressure point. C) X-axis profiles at Z = 0.1mm, the height of the cells in the well.

### Acoustic Intensity, Pressure, and Thermal Effects

Acoustic intensity variations may impact the distribution of acoustic pressure distribution due to non-linearities in absorption and scattering during sonication. Changes to the distribution of the ultrasound field at varying acoustic intensities could impact the ultrasound dose delivered to cells between setups. However, at low acoustic intensities, such as those intended for the SDT proposed in this paper, the impact of this effect not well established ^30^. Therefore, systematic studies to assess the pressure field distributions at varying acoustic intensities were also performed.

In-well submerged FUS field mappings were carried out at varying acoustic intensities of 0.4-1.6W/cm^2^ with the water level set to 60mm above the 75mm focus point of a 666kHz transducer. The mapped peak pressure at each intensity shows a progressive increase in the peak pressure within the well (100-200kPa) (Figs 4 Ai-Di). Comparison of non-standardised-scale pressure maps (Figs 4 Aii-Dii) showed a consistent pressure field with increasing intensity, demonstrating that these intensities are sufficiently low to avoid disruption of the field. Therefore, the target pressure and cumulative sonication dose can be varied between wells within a pre-programmed experiment for a rapid testing of parameter configurations.

**Figure 4.**
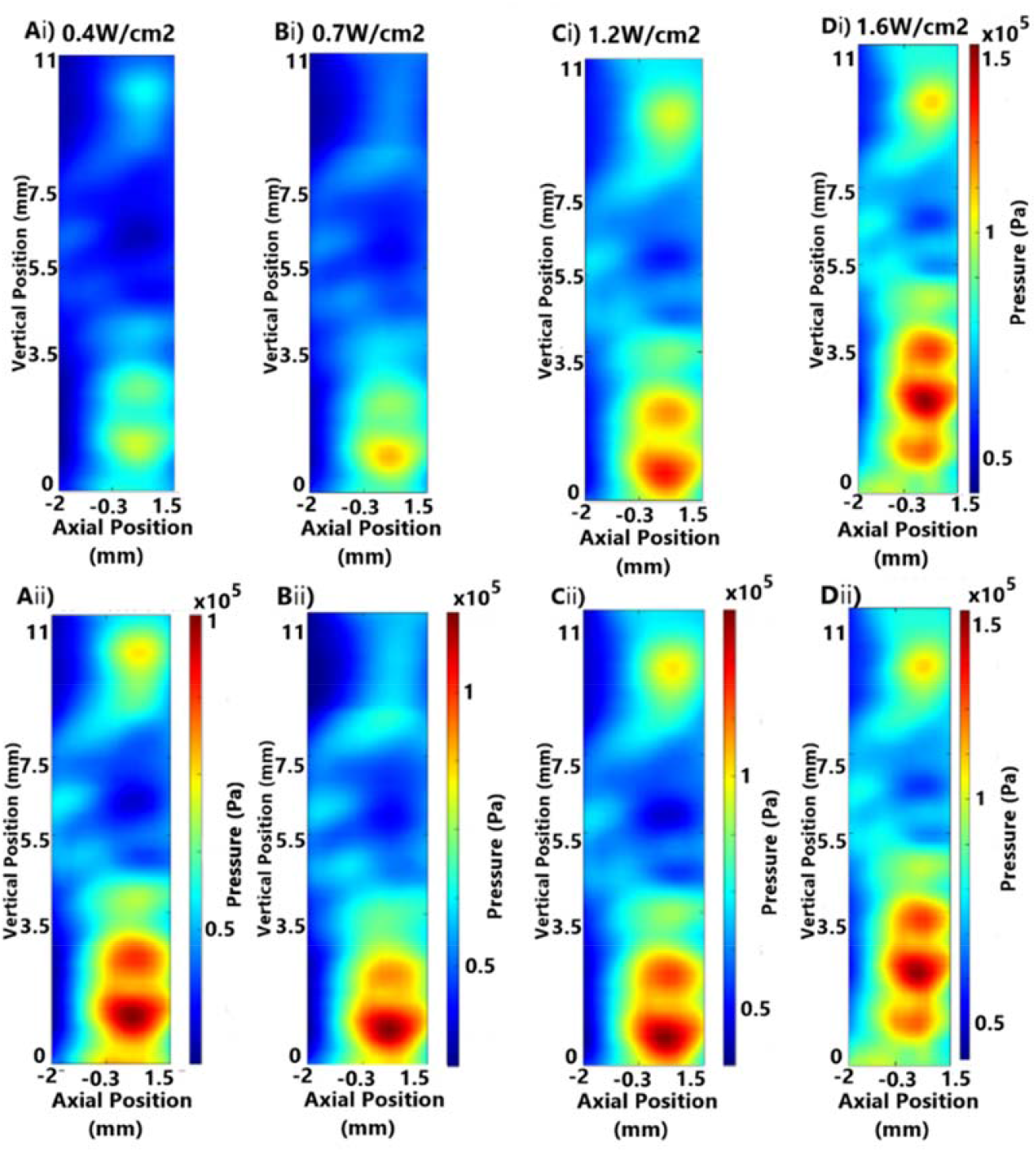
Comparison of in-well (submerged) pressure field mappings at increasing emitted intensity, emitted generator amplitude, and target peak pressure of continuous wave focused ultrasound, with and without standardised scale bars (i and ii subfigures, respectively): Ai and Aii) 5.4W/cm ^2^ (2.5V, 0.402MPa) Bi and Bii) 7.0W/cm ^2^ (2.75V, 0.458MPa), Ci and Cii) 9.9W/cm ^2^ (3.25V, 0.545MPa) Di and Dii) 12.2W/cm2^2^(3.5V, 0.603MPa).

Further, we also investigated the occurrence of thermal effects due to FUS exposure. High frequency focused ultrasound can carry a significant thermal dose dependant on acoustic intensity and may cause target cell hyperthermia. Regardless of low-intensity ultrasound parameters utilised (<15w/cm ^2^, <15% duty cycle), no thermal dose was detectable at the end of a 5-minute sonication. The difference between the target well and average temperature of the 8 surrounding wells did not exceed 0.3°C. Further information is available in the supporting information - *thermal effect* (Supplementary Fig S4).

### Standing Waves and Vertical Interference

In our system, ultrasound waves meeting the fluid-air boundary at the water’s surface reflect downwards and may interfere with ascending waves to produce standing waves (a phenomenon that is well described in the literature). The degree of ultrasound interference, and thus potential for standing waves, is dependent on the transducer’s curvature and the distance from the focus point to the fluid-air boundary, as the intensity of reflected waves will reduce in line with the inverse square law. Focused ultrasound transducers typically utilise a concave surface to allow the waves to focus to a point; as such a degree of lateral dispersion may be seen beyond the focus point. Therefore, standing waves may be mitigated in submerged experimental setups.

Comparison of the degree of ultrasound field distortion between in-well (water level) and in-well (submerged) setups was undertaken (Fig 2). The distance between pressure field oscillations increased from 0.5mm to 1.2mm when the plate was submerged (666kHz transducer), indicating a more consistent pressure field. The diameter of the lowest peak node, indicating the area of cells to which peak pressures have been applied, remained constant however, the in-well (submerged) setups showed a greater intensity at the base of the well while the in-well (water level) setups showed a lower, but more consistent central cell exposure (0.3MPa compared to 0.25MPa). The point of peak intensity was raised in the in-well (water level) setup from the base of the well by 2.5mm.

Further, the effect of lateral acoustic boundaries across the wells where multi-well microplates showed varying resonant properties; adjacent wells containing fluid are relatively hypo-resonant, and the air-filled inter-well spaces are relatively hyper-resonant. This inconsistent ‘lateral dampening’ effect was exacerbated at unequal fluid levels between adjacent wells (Supplementary Fig S5).

It is therefore prudent to be restrictive when sampling: beyond a radius of ∼0.9mm, cells do not receive a comparable dose to those within that boundary. The validity of this method is corroborated by the analysis of the microclear plate x-lines, see *‘plate base material and thickness,’* above. Additional measures to mitigate standing waves, including PVA inserts, are discussed in the supporting information (Supplementary Fig S6).

### Frequency

Frequency impacts several factors in in-vitro SDT setups. With higher frequency transducers, the focus distance is extended, and the field profile is narrower (Figures 5A & 5C). Consequently, the reflected FUS from the fluid-air boundary above ‘doses’ the target well with a higher intensity at higher frequencies (assuming an equivalent submersion depth). Acoustic pressure simulations confirmed the presence of the reflected waves, clearly illustrated in the 1.2MHz open field mapping (Fig 5C). Changes to frequency would alter the severity of standing wave reflections during the in-well testing with the cells at the water level. 666kHz and 1200kHz transducers were compared in 2 settings via the use of pressure field simulations (Fig 5). For ‘no well plate’ and in-well (submerged) setups, the lower frequency transducer showed a reduction in the frequency of pressure field oscillation and improved consistency for the estimation of FUS dose. In addition to this, the FUS field is broader for the 666kHz transducer compared to that of the 1.2MHz transducer. The benefit of a broader field is that the usable cell sample area is expanded owing to a more consistent intensity across the width of the well as described above.

**Figure 5.**
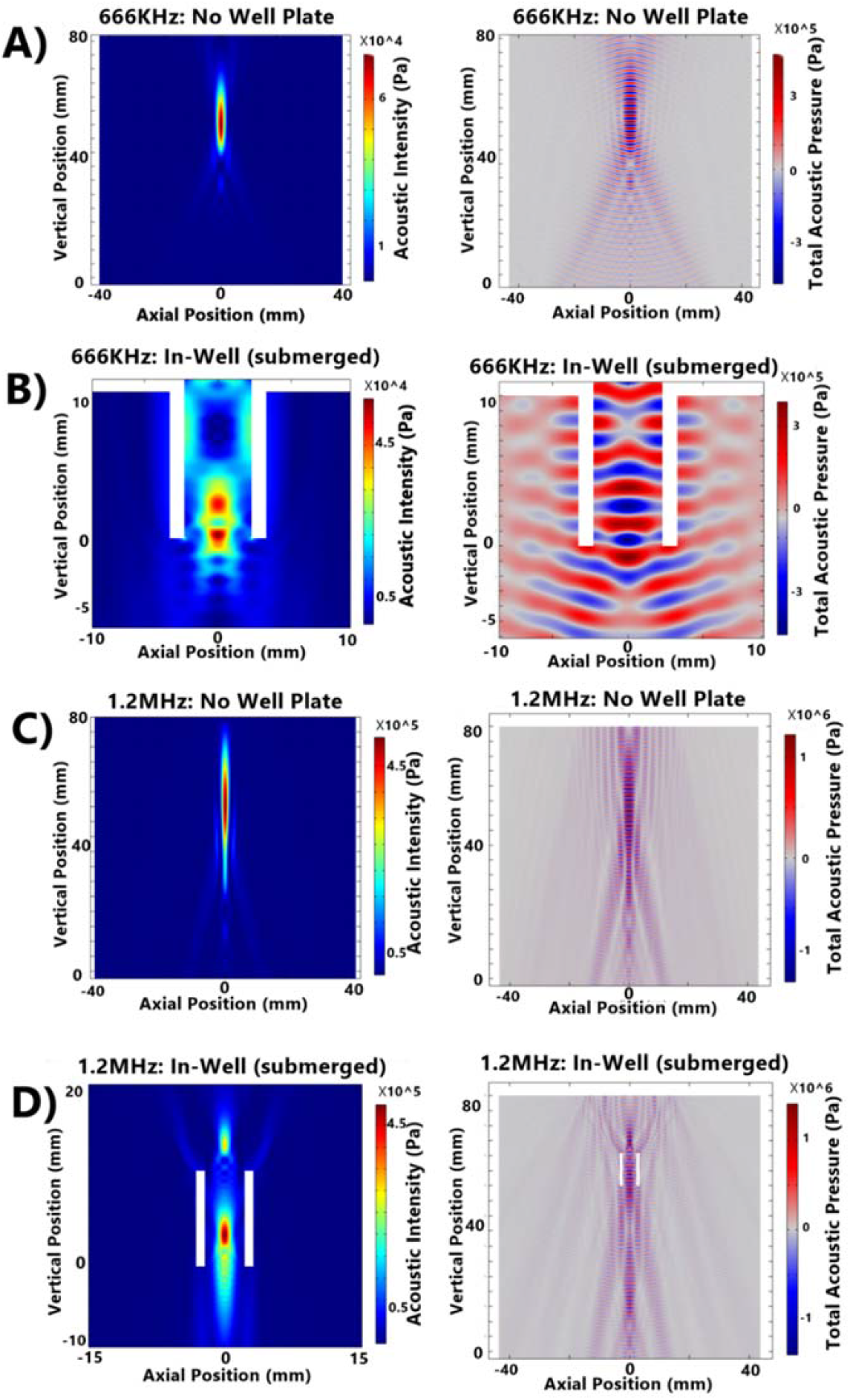
COMSOL Simulations of in-well (submerged) and no well plate setups for 666kHz and 1.2MHz transducers. Acoustic intensity and total acoustic pressure are shown for each condition: A) 666kHz No Well Plate Setup, B) 666kHz In-Well (Submerged) Setup, C) 1.2MHz No Well Plate Setup, D) 1.2MHz In-Well (Submerged) Setup

Axial pressure profile of 666kHz and 1.2MHz transducers were compared to assess field variability at the level of their respective peak pressures and at the base of the well. While both frequencies demonstrated oscillations throughout, the amplitude of 1.2MHz field was observed to be lower (supplementary Fig S5). The 1.2MHz showed a lower coefficient of variation of 29.9% owing to a taller central peak compared to 666kHz which held 42.9% due to more predominant oscillations. Further, higher frequency was quantifiably more consistently across the vertical axis. Figure 5B which depicts the profile of the field at the level of the cell-lines in the well and the profiles, were observed to be similar.

### SDT impedes glioma cell growth

A previous study has utilised U87MG and U251 cells to establish SDT parameters ^31^. However, the validity of both U87MG and U251 cells as glioma models have come under serious scrutiny ^32 33^. Hence to establish the effect of SDT on state-of-the-art glioma cells, primary patient derived cells GBM22 and glioma stem neurospheres GBM120 were utilised. Upon SDT exposure, GBM22 cells exhibited marked cell death (Fig 6A) along with enhanced Annexin V apoptotic signal (Fig 6B & Supplementary Fig S7). The cell death phenotype was not observed upon individual exposure to photosensitiser 5-ALA or focused ultrasound alone suggesting the combination of 5-ALA and FUS is required to induce apoptosis. Interestingly, apoptotic signal was exclusively present in the SDT samples with proportional increase observed with increasing dose of FUS (Supplementary Fig S7). Furthermore, SDT exposure led to increased reactive oxygen species induction as evidenced by positive signal with CellROX (Fig 6C). SDT induced ROS upregulation in 100% of attached live GBM22 cells (Fig 6C).

**Figure 6.**
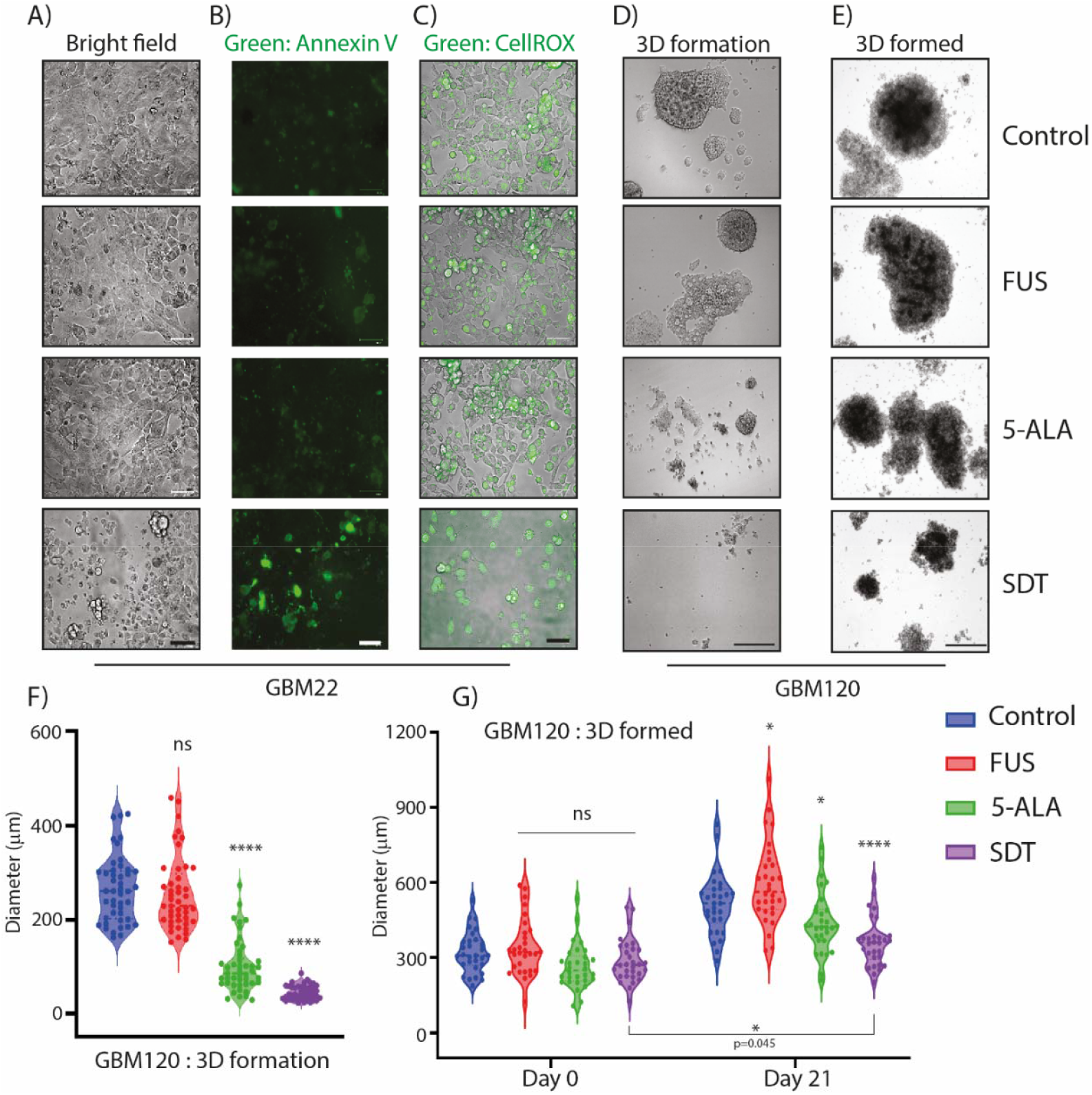
SDT induces cell death in primary patient derived glioma cells in vitro. A) Representative bright field images of GBM22 cells 1.5h post-treatment with/without 1mM 5-ALA, with/without FUS (0.4w/cm^2^, 30s cumulative sonication, 10% DC, 90ms pulse length), and both (SDT). Scale bar= 125μm. B) Representative fluorescent images only of GBM22 cells, pre-treated with Annexin V-FITC apoptosis marker, 1.5h post-treatment with/without 1mM 5-ALA, with/without FUS (0.4w/cm^2^, 60s cumulative sonication, 10% DC, 90ms pulse length), and both (SDT). Scale bar= 125μm. Also see Supplementary Fig S7. C) Representative overlaid bright field and fluorescent images of GBM22 cells, pre-treated with CellROX dye, 1.5h post-treatment with/without 1mM 5-ALA, with/without FUS (0.4w/cm^2^, 30s cumulative sonication, 10% DC, 90ms pulse length), and both (SDT). Scale bar= 125μm. D) Representative bright field images of GBM120 3D neurospheres. The dissociated GBM120 cells were pre-treated with indicated conditions as in A and allowed to form neurospheres over 21 days. n=3 biological replicates. Scale bar= 200μm. E) Representative bright field images of GBM120 3D neurospheres. The indicated treatments as in A were carried out on pre-formed 3D neurospheres and allowed to grow over a further 21 days. n=3 biological replicates. Scale bar= 200μm. F) Quantification of D across all n=3 replicates. The diameter of the neurospheres were quantified using ImageJ. The significance of the differences was measured using one-way ANOVA with Dunnett’s multiple comparisons. ****p < 0.0001; ns: not significant. G) Quantification of the diameter of formed neurospheres on Day 0 and Day 21 across all n=3 replicates. The diameter of the neurospheres were quantified using ImageJ. The significance of the differences was measured using two-way ANOVA with Bonferroni’s multiple comparisons. ****p < 0.0001; *p < 0.05; ns: not significant.

To understand if SDT exposure can target 3D glioma stem neurospheres, GBM120 cells were treated with or without 1mM 5-ALA, FUS, and SDT. The cells were then cultured over 21 days and allowed to form 3D neurospheres in stem media. Interestingly, the SDT exposed population of GBM120 cells exhibited the smallest 3D spheroids compared to the 3 controls. 1mM 5-ALA alone did exhibit some toxicity to GBM120 cells, but SDT treatment was the most effective in reducing glioma stem neurosphere growth (Fig 6D&F). Similar results were obtained when GBM120 pre-formed 3D neurospheres were exposed to SDT (Fig 6E&G). 5-ALA alone toxicity was modest but significant in the 21-day cohort and SDT treatment remarkably affected 3D spheroid growth progression over 21 days. SDT-treated cohort did show a very modest yet statistically significant growth over 21 days suggesting incomplete cell kill (Fig 6G). Cells lysed 1.5hrs post treatment showed an increase in phospho-ERK signal in 5-ALA, FUS, and SDT treatment groups compared to control while a very modest increase in phospho-AKT Ser473 and Thr308 were observed in 5-ALA treatment alone (Supplementary Fig S8). Poly-D-lysine used to ensure immobilization of neurospheres for sonication altered the distribution of growth patterns from spheroids to more irregular patterns in some neurospheres (Fig 6D&E). This prevented the accurate calculation of volume; hence diameter was used as a simplified measure and a greater sample size was collected to mitigate the effects of irregular growth. Since the centre of the well saw the highest peak pressure and acoustic intensity compared to well periphery (Fig 3C), neurosphere sampling and imaging was conducted from the centre outwards, so that the results reflected the most consistent and predictable FUS dose parameters available within the current system. Neurosphere sampling was conducted via averaging 2 perpendicular diameters of the 5 neurospheres closest to the centre of each well, the strongest point of the FUS peak pressure map. A total of 45 neurospheres were measured per condition every 3 days for 21 days over 3 biological replicates with 3 technical replicates in each.

## Discussion

An accurate determination of FUS dose and consistency in its application to cells is crucial to the translation of in vitro SDT experiments. We have designed our system for this purpose and sought to optimise field mapping to later correlate to cellular effects, rather than to investigate one physical property as is often reported in literature. Further studies should be undertaken to refine similar experimental setups and tumour models for the purpose of rapid testing and optimisation of sonication parameters. With further understanding of the physical and cellular mechanisms involved, in vitro data may inform the evolution of ideal human trial FUS parameters, a significant factor in the clinical application of SDT.

Limited data exists on the impact of SDT in inducing cell apoptosis. For this reason, standard parameters for cell testing were derived from the most efficacious single-transducer pulsed-FUS parameters in rats ^24^. A lower acoustic intensity was utilised to facilitate testing (table 1), however, duty cycle, pulse length, and cumulative sonication dose were broadly maintained. The impact of both rat skulls and standard 96-well plates will reduce the target FUS intensity, though the degree may vary. It is important that in vitro parameter selection is realistic when translated to patients. No thermal effect was seen at the selected intensity; alteration to duty cycle may have implications on skull heating effects in vivo. There is an inverse relationship between duty cycle and to complete a set cumulative sonication. Patient may not tolerate treatment of over 90-100 minutes, for 18 points to be sonicated as in Wu et al, 2021, a 10% duty cycle was selected to apply 30s of cumulative sonication to each point in this treatment duration.

**Table 1.** Comparison of focused ultrasound parameters (acoustic intensity, pulse length, duty cycle, and cumulative sonication time) used by Wu et al (2021) to those included in this study.

| Study | Acoustic Intensity (w/cm<sup>2</sup>) | Pulse length (ms) | Duty cycle (%) | Cumulative Sonication (seconds) |
| --- | --- | --- | --- | --- |
| This paper | 0.4 | 90 | 10 | 30 |
| Wu et al (2021) | 5.5 | 86 | 8.6 | 24 |

The exact mechanism of PpIX activation and wider cellular impact of focused ultrasound is yet to be fully described. Further development of in vitro systems will allow the investigation of these mechanisms. This system may be used to examine the cellular effects of varied focused ultrasound parameters in comparison to in vivo trials when standardised for the level of PpIX activation. The use of this surrogate marker for cavitation-induced sonoluminescence accounts for possible differences in ultrasound dynamics in vivo, for instance the impact of circulating oxygen. Though this system cannot currently examine the physical mechanism of sonodynamic therapy, it provides a strong basis for studies into the cellular mechanism and its efficacy.

The mitigation of standing waves is one of the key factors within FUS experimental setups. Standing waves create consistent nodes and anti-nodes which may have amplified effects; however, these are predictable and can be measured. Sampling cells from known nodes allows for estimation of pressures they have been exposed to in setups in which standing waves have not yet been fully eliminated. The further development of cell-safe methods to alleviate standing waves, including PVA inserts, may improve field consistency.

The needle hydrophone used to generate field maps required a minimum of 400μL within each well, to prevent damage to the equipment. Most cells tests are undertaken with 100μL of media in each well, which is expected to cause further standing wave reflection in simulations. The X and Z axis movements of the hydrophone may have disrupted the water’s surface causing ultrasound scattering, as such, the interval between movement and hydrophone detection was increased. The comparison of more complex setups to simulations may improve dose estimations where measurement via a hydrophone is not possible, including sealed and submerged cell enclosures.

Furthermore, 5-ALA treatment activated the proliferative AKT and ERK signalling (Supplementary Fig S7). This was expected as 5-ALA is an amino acid derivative and potentially a nutrient. The reduced growth phenotype in 5-ALA alone cohort may be due to toxic metabolites accumulation over time in the cell media (Fig 7D-G). This toxicity may not be an issue in patients as the exogenous amino acid is excreted out quickly and is FDA-approved for ingestion. Interestingly, only phosphorylated ERK was observed upon FUS treatment (Supplementary Fig S7) which could be a response to the stress. While FUS alone stress is potentially non-fatal, the combination of 5-ALA and FUS is indeed highly toxic and tips the scales for glioma (Fig 7). This apoptotic cell death observed in SDT alone could be partly due to excess ROS production. The slight increase in FUS–alone cohort over 21 days (Fig 7G) is probably due to a broader distribution of initial diameter of neurospheres. The SDT treated cohort also exhibited a modest growth over 21 days suggesting that some cells remain proliferative within the neurospheres. It would be interesting to check if combination with chemotherapy could alleviate this modest growth.

An understanding of factors influencing ultrasound fields and the mitigation of interference is necessary for the accurate determination of cell dosing in in vitro studies and the translatability to clinical sonodynamic therapy. Herein, we report an optimised system that can induce death in glioma 2D and 3D glioma cells in vitro and sets the basis for further understanding of SDT and its therapeutic potentials in cancer.

## Additional information

### Author contribution

AK, TM, ZG, and HH collected the data included in this paper. LGN, AS, JN, HL, LL contributed to resources and acquired the data. SB and JJ Supervised the project. LL, AM, KHI, ZH, SB, and JJ designed the experiments and advised on the acquisition of data. All the authors contributed to the manuscript preparation.

### Data availability

All data that support these findings of the present study are included in the manuscript and supplementary information. Further information is available upon request to the corresponding author, James Joseph.

## Appendix

**1.x**

| Item Number | Name | Manufacturer |
| --- | --- | --- |
| 1.1 | T08200 ETH 300W | Hydor, Vicenza, Italy |
| 1.2 | LET 775, 12V, 65PSI | Lei Te Co., China |
| 1.3 | 1.0mm Needle Hydrophone | Precision Acoustics Ltd, Dorchester, UK |
| 1.4 | PicoScope® 4262, 16-bit high-resolution oscilloscope | Pico Technology, St Neots, Cambridgeshire, UK |
| 1.5 | Standard clear plate: Nunc™ MicroWell™, Nunclon Delta-Treated, Flat-Bottom Microplate | ThermoFisher Scientific, Waltham, Massachusetts, United States |
| 1.6 | Ultrasound-translucent plate: µClear® (190µm film-based), Black polystyrene constructed chimney-well, tissue-culture treated, sterile, 96-well plate with a polystyrene lid. | Greiner Bio-One, Stonehouse, England, UK |
| 1.7 | Motorized XSlide™, XN10 Series, Lead Screw | Velmex Inc., New York, USA |
| 1.8 | VXM™-2 Motor Controller | Velmex Inc., New York, USA |
| 1.9 | thermoIMAGER TIM 160 | Micro-Epsilon UK Ltd., Liverpool, UK |
| 1.10 | 666kHz ultrasound transducer | Precision Acoustics Ltd, Dorchester, UK |
| 1.11 | 1193kHz ultrasound transducer | Homemade in the lab |
| 1.12 | AFG3101, Single Channel, Arbitrary/Function Generator | Tektronix UK Ltd., Bracknell, Berkshire, UK |
| 1.13 | Poly(vinyl alcohol), 99.3-100.0% hydrolysed, M.W. approx. 146,000-186,000, ACROS Organics™ | Fisher Scientific UK, Loughborough, UK |

