## Supplementary info for "Development and Optimisation of in vitro Sonodynamic Therapy for Glioblastoma"

### Supplementary methods:

*System Control and Calibration:*

Multiple NH field mapping setups were developed, diagrams of these are shown in Figure S1.

The system must be calibrated to ensure accurate FUS dose estimation and concordance to pressure field simulations. The system can be calibrated manually, where the hydrophone is adjusted in X, Y, and Z axis to find the focal point and the base of the target well is adjusted to this point. Automatic calibration, whereby the focal point was determined in the same way, however X and Y axis limit switches position the well plate. As long as the transducer position remains constant, this method allows for accelerated testing protocols. Peak pressure profiles to compare manual and automatic calibration were performed (figure S1), though slight X axis deviation was seen, there was negligible difference in peak pressure (121kPa with continuous wave 0.4w/cm2). The FUS focal point was always within the target well. A reset command was developed to allow the recalibration of the system from any position, this gave an error of 1 to 2mm, as such it was supplemented by fine manual calibration (Fig S2). The diameter of the focus point may be altered by transducer frequency and curvature of the transducer surface. When a 666kHz transducer is utilised, the focus point diameter of the 666kHz transducer is 8mm, roughly equal to the inner diameter of a well of a 96-well plate. The effects on adjacent wells is minimal. (Fig S3).

*In well (PVA – Submerged):*

The gel preparation prior to casting was completed in line with previous literature (1). A 10% solution of PVA (Poly(vinyl alcohol), 99.3-100.0% hydrolysed, M.W. approx. 146,000-186,000, ACROS Organics™, Fisher Scientific UK). was prepared using sterile, degassed, and de-ionised water (manuscript appendix 1.12) and set into a standard polystyrene 96-well plate prior to freeze-thawing to establish internal structure. The cast was then divided into manageable inserts of 3 by 3 wells for field testing, inter-well spaces were represented in the cast and included in the PVA gel. A 2mm diameter tunnel was made vertically through the central axis of the central well of the gel and plate lid to allow the NH to be inserted. The central well received 40µL of water and 40µL of PBS was placed into the 8 surrounding wells. Inter-well spaces received 18µL of PBS, equivalent in fluid level to the surrounding wells. The insert displaces the fluid within each well around the column of PVA descending into each well, eliminating the fluid-air boundary below the PVA. PBS was placed on the PVA to eliminate an air pocket underneath the plate lid. The plate was sealed by its insertion into the plate holder before being submerged 60mm under the surface of the water bath. To map the field within the PVA material, a pinhole was created in the centre of one well's cast, allowing the tip of the hydrophone to be inserted into the well. In this method the movement of the probe was constrained to the Y axis. The results were plotted as a line graph of pressure readings against the height above the floor of the well.

### Supplementary Results:

*Thermal effect:*

To determine the thermal effect of low intensity ultrasound, mock cell tests were conducted at the higher and lower ranges of acoustic intensity and duty cycle used in low-intensity SDT experiments. 100µL GBM media with 100µL PBS border was sonicated, with the fluid level set to the focus distance, using the following parameters and recorded under thermal camera (Fig S4A and B): 0.4W/cm2 (0.1MPa) and 2W/cm2, 50ms pulse (5% duty cycle) and 150ms pulse (15% duty cycle). For all tests, the pulse frequency remained at 1Hz with a transducer frequency of 666kHz. All target wells received 30 seconds of cumulative sonication during which continuous monitoring of the average temperature of both the target and target with surrounding wells was recorded and graphed (figure S4C, D, E, and F). The temperature remained constant with change remaining under 0.3 degrees Celsius. These findings are in line with literature (1), in which temperature rise is equivalent to 0.15 degrees per 1W/cm2 of continuous wave sonication. Our results show that as we use pules with a duty cycle of 15% or less, we see no significant thermal rise. If continuous wave, from literature data we may estimate a 0.32 degree rise from pre-post sonication thermal data.

*Lateral Dampening and Adjacent well interactions*

Observable patterns were produced in the ultrasound field within the target well. Viewed from above, relative ultrasound dampening at 90° increments was seen, in line with fluid-filled wells, and relative ultrasound reflection offset at 45° intervals, in line with the empty inter-well's spaces. This was also observed in the vertical axis where the height of the fluid level varied between the wells. In NH field mapping, when the target well is filled with the required 400µl of fluid, and adjacent wells contain just 100µl of fluid, as shown in figure 2, there is a noticeable region of reduced pressure, roughly rectangular in shape, extending inwards from the outside edge of the well by approximately 2mm, and upwards by approximately 1mm, corresponding to the height of the fluid in the adjacent well and the boundary between hypo-resonant fluid and hyper-resonant air space. This effect is referred to herein as 'lateral dampening'.

The lateral dampening effect was also assessed using the two axial profiles taken from the ultrasound field within a well at 0.3mm and 2.3mm above the base. As shown in Figure 5A, lateral dampening was observed with 100uL of fluid in the adjacent wells. The relative width of the *peaks* of each wave were comparable (taken to be the region of the curve greater than the mean), however, the curve taken at a height of 0.3mm rapidly attenuated beyond a radius of 0.9mm. That was not seen in the curve at 2.3mm height where the field is sustained above -1 standard deviation until a radius of 1.9mm. Further, the attenuation of the field was demonstrated using serial z-lines taken at regular intervals from the central axis (Supplementary Fig S5B). As expected, above a height of 1.6mm in the well, the pressure diminished relatively evenly with an increase in the radius (axial distance). As the plot approaches the base of the well (decreasing height), the two innermost radii maintained a regular field pattern. In contrast, the plots at the two outermost radii (and to a lesser extent at a radius of 1.1mm) were flattened, demonstrating dampening at lower and more lateral points in the well.

*Frequency*

Frequency impacts several factors in in-vitro SDT setups. With higher frequency transducers, the focus distance is extended and so the submersion depth for a given setup as a proportion of focus distance is reduced. This causes a relative reduction in scattering and subsequent increase to vertical interference returning to the target well, as r is defined as the focus distance in estimations using the inverse square law. The focus length of the 666kHz and 1200kHz transducers, described in this paper, were 75 and 125mm, respectively.

*PVA inserts and a continuous fluid column:*

The inclusion of specialised PVA cryogel inserts may provide an opportunity to reduce the reflected ultrasound dose. Firstly, if placed beyond the target cells, by absorbing ultrasound there is less available dose to return to the cells. Secondly, where inserts are used to eliminate air pockets between the surface of cell media and the underside of the plate lid, a continuous fluid column absent of fluid-air interfaces can be established. A plate may then be temporarily sealed and submerged in the water bath to allow ultrasound to propagate beyond the plate to the surface of the water bath, dissipating the intensity of the returning waves.

Z-line pressure profiles of this setup are similar to that of a standard polystyrene plate submerged at an equal distance (Supplementary Fig S6). Both profiles show a peak pressure from 1.9-3.3mm above the base of the well and demonstrates the lack of PVA interference in in-vitro experimental setups. PVA can absorb ultrasound waves, preventing their return and interference, this is demonstrated by a reduction in both peak and well-base pressures by 60% and 80% respectively on the introduction of a PVA insert (0.25MPa to 0.10MPa and 0.14 to 0.03MPa). This system with optimisation is a potential method for the mitigation of vertical dose reflection in cell line testing.

Sterilised PVA gels made with degassed water may provide a temporary solution during testing, however, testing of the cell tolerability and impact on 3D cultures should be completed prior to use. Sealing the plate prior to submersion may pose the possible concern of cell hypoxia in long, automated plate testing setups. As the interactions between hypoxia, dissolved gases, microbubbles, focused ultrasound, and the effects of ROS are not fully understood, this may introduce compounding factors into experimental designs and should be carefully considered.

**Figures**

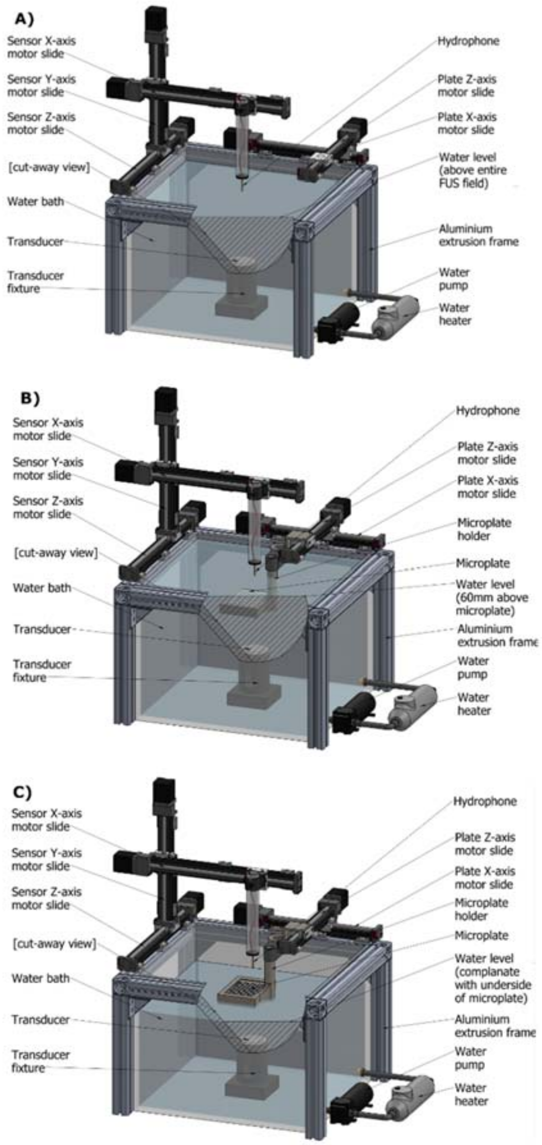


**Figure S1** – Diagram of NH field mapping setups as described in *methods and materials*;

A) *No well plate:* The well plate was removed from the testing setup to allow the open field to be mapped within the water bath, with the water level well above the focal point of the US field. The focal point was identified by manually jogging the hydrophone until the peak sound pressure was located. This also served as a reference point for the centre of the mapped area.

B) *In well (water level):* The well plate and thermal camera were included in the testing setup and the stage was adjusted to place the well plate at the level of the water surface and with the focal point of the ultrasound within the target well. The thermal camera was focussed on the target well which was filled with 400μL water. Surrounding wells contained 100μL of phosphate buffered solution (PBS).

C) *In well (submerged):* The well plate was included in the testing setup and the stage was adjusted to place the well plate 60mm below the water surface and with the focal point of the ultrasound within the target well.

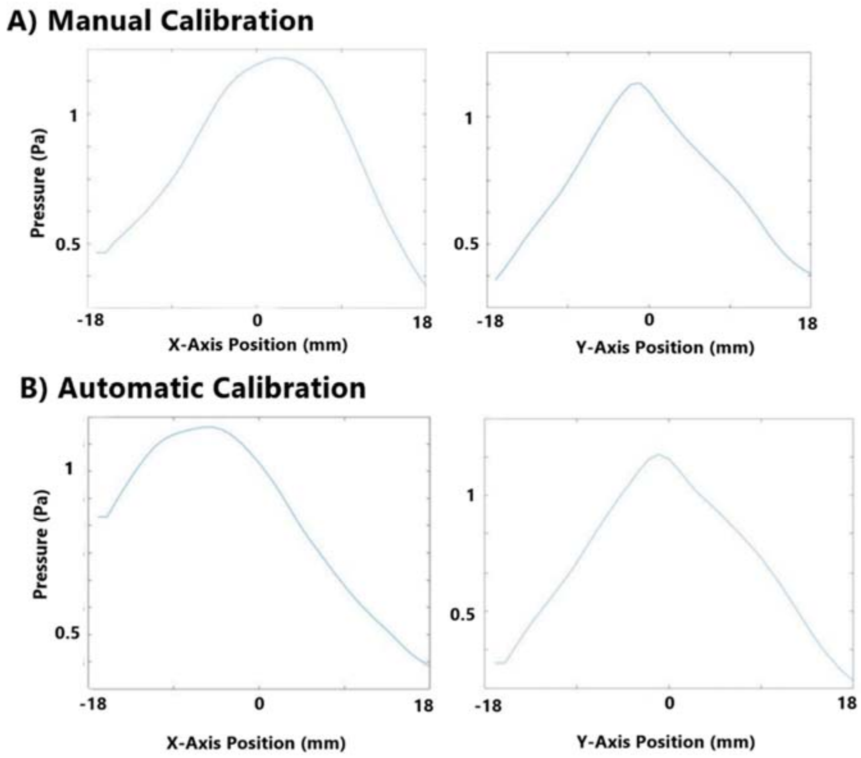


**Figure S2** - X and Y axis peak-peak pressure profiles for manual and automatic calibration of the system.

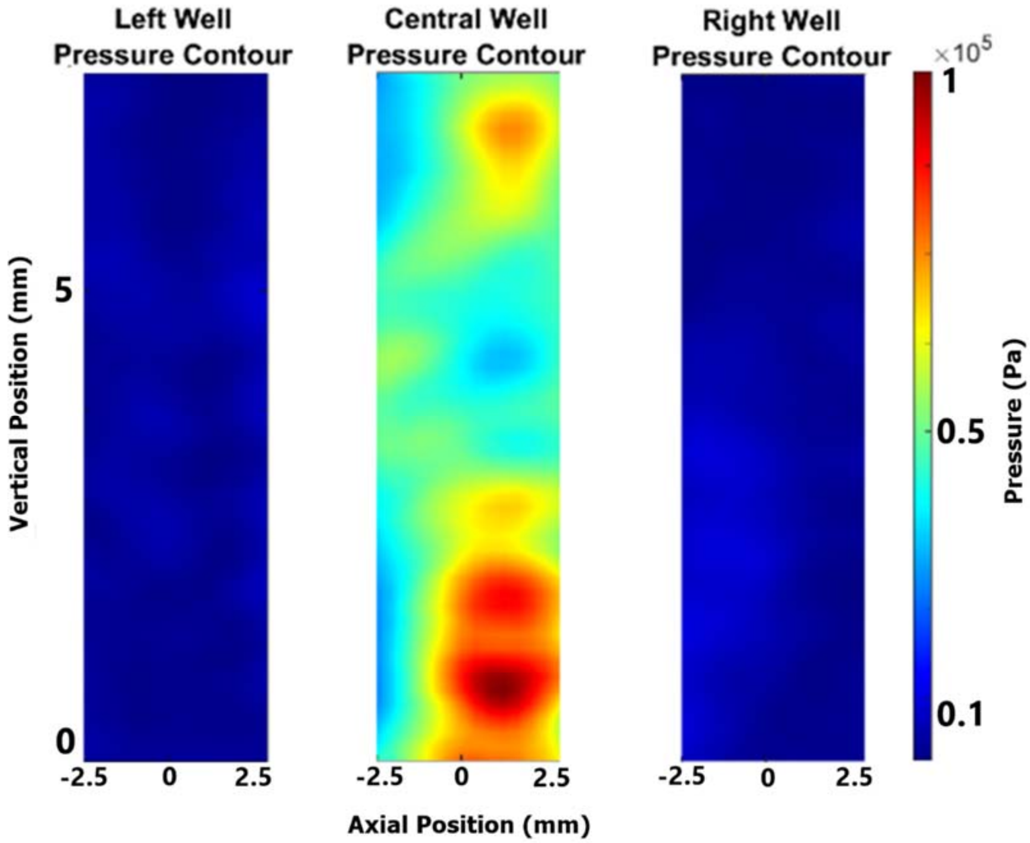


**Figure S3** – Standardised scale peak-peak voltage mapping, peak-peak pressure mapping, and normalised decibel mapping for target wells and adjacent wells to the left and right of a calibrated target.

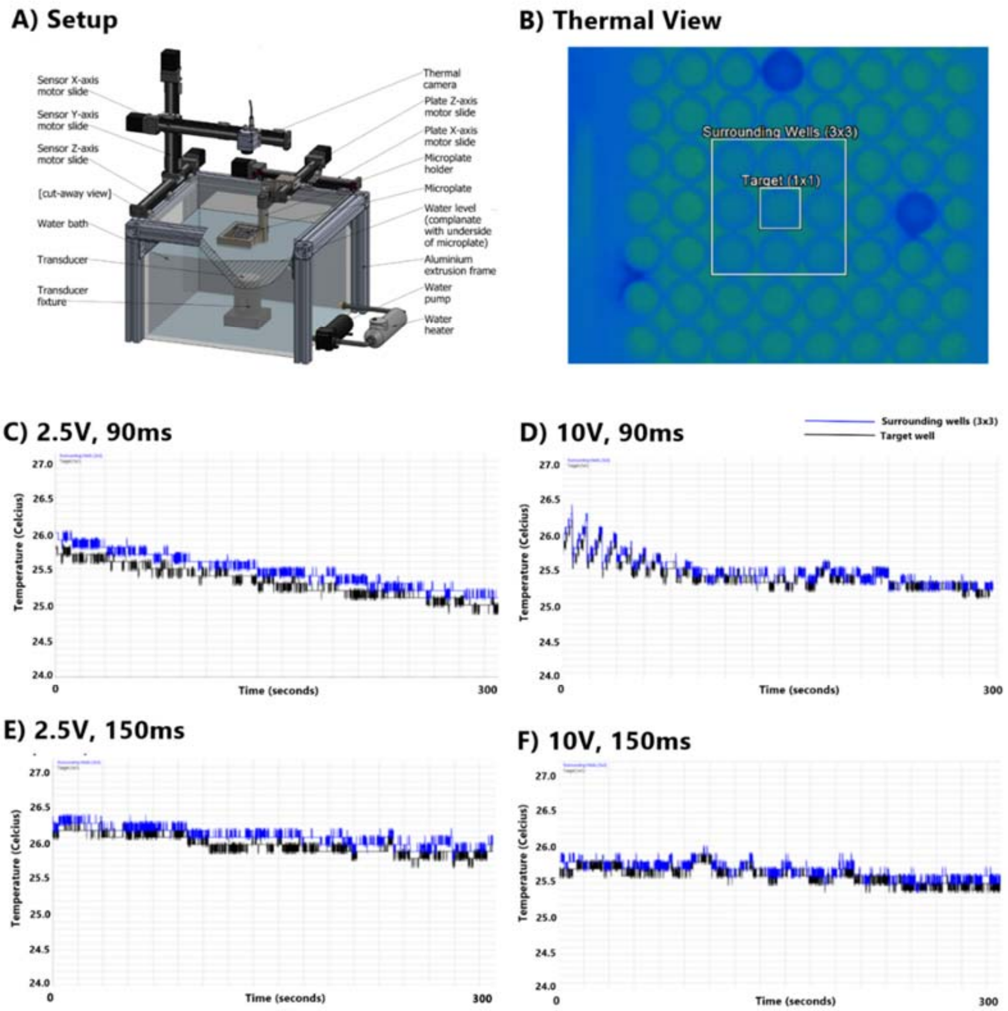


**Figure S4** – Thermal effect imaging at varied sonication parameters:

A) Diagram of in vitro experimental setup for the determination of thermal effects (in-well (water level) setup with thermal camera aligned with central axis of the FUS transducer).

B) Thermal camera images taken at the end of sonication of mock cell tests under varied intensity and sonication parameters. Lines over time together as separate combined figure. Scale bar with peak temperature of 15% DC at 2W/cm2.

C) Temperature time graph for 2.5V generator amplitude with 90ms pulse length (9% duty cycle). Average temperature of measure areas of the target well (black) and target well with 8 surrounding wells (blue) for in-well (surface level).

D) Temperature time graph for 10V generator amplitude with 90ms pulse length (9% duty cycle). Average temperature of measure areas of the target well (black) and target well with 8 surrounding wells (blue) for in-well (surface level).

E) Temperature time graph for 2.5V generator amplitude with 150ms pulse length (15% duty cycle). Average temperature of measure areas of the target well (black) and target well with 8 surrounding wells (blue) for in-well (surface level).

F) Temperature time graph for 10V generator amplitude with 150ms pulse length (15% duty cycle) - Greatest intensity. Average temperature of measure areas of the target well (black) and target well with 8 surrounding wells (blue) for in-well (surface level).

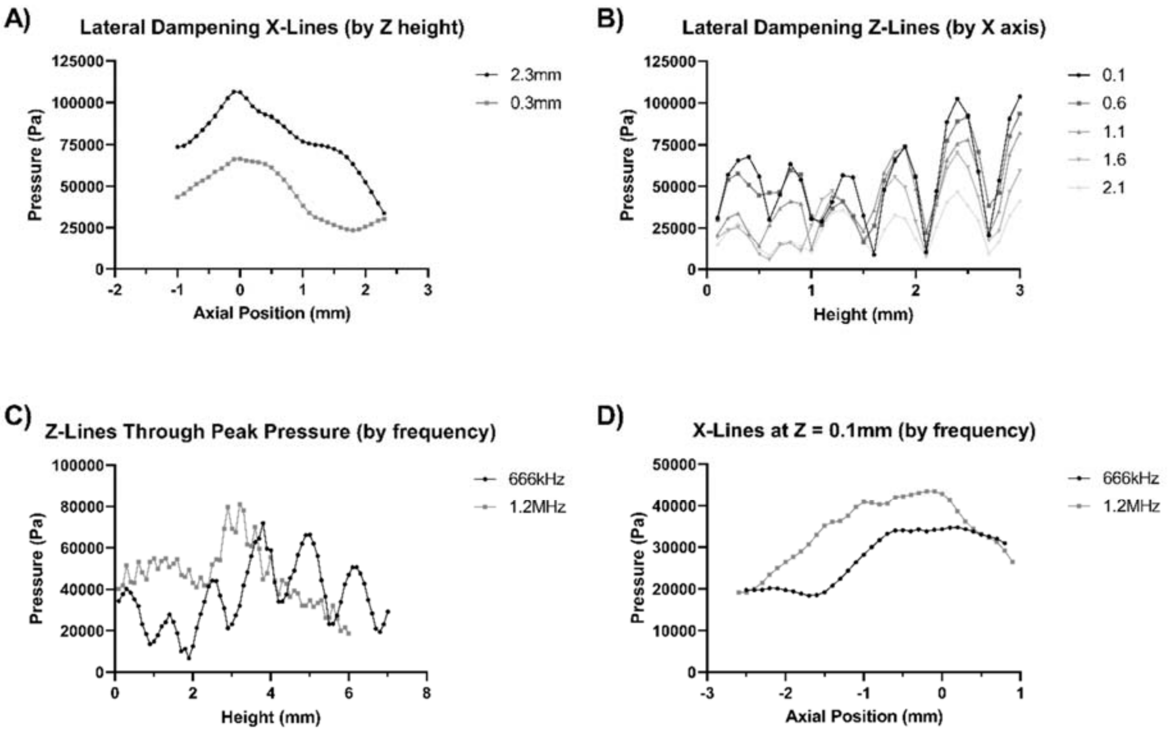


**Figure S5** – X and Z Pressure profile comparisons illustrating the effects of altered frequency and adjacent structures on pressure distribution

A) X-axis pressure profiles comparing lateral dampening from below and above the water level of adjacent wells, 0.3mm and 2.3mm high, respectively, in an in-well (water level) setup.

B) Z-axis pressure profiles at 0.1, 0.6, 1.1, 1.6, and 2.1mm from well central axis for an in-well (water level) setup.

C) Comparisons of 666kHz and 1.2MHz pressure profiles for 0.4W/cm$^2$ continuous wave focused ultrasound in an in-well (submerged) setup for Z-axis profiles intercepting the peak pressure point.

D) Comparisons of 666kHz and 1.2MHz pressure profiles for 0.4W/cm$^2$ continuous wave focused ultrasound in an in-well (submerged) setup for X-axis lines at Z = 0.1mm, the height of the cells in the well.

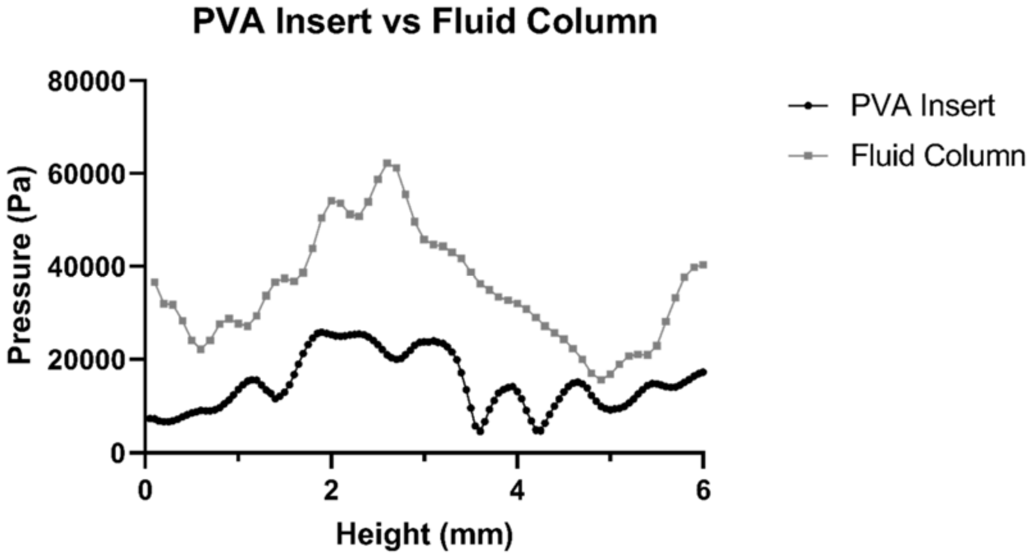


**Figure S6**– Submerged plate testing of Z-Line pressure profiles between a plate containing a PVA insert and a plate without inserts or a plate lid (equivalent to a perfect fluid column).

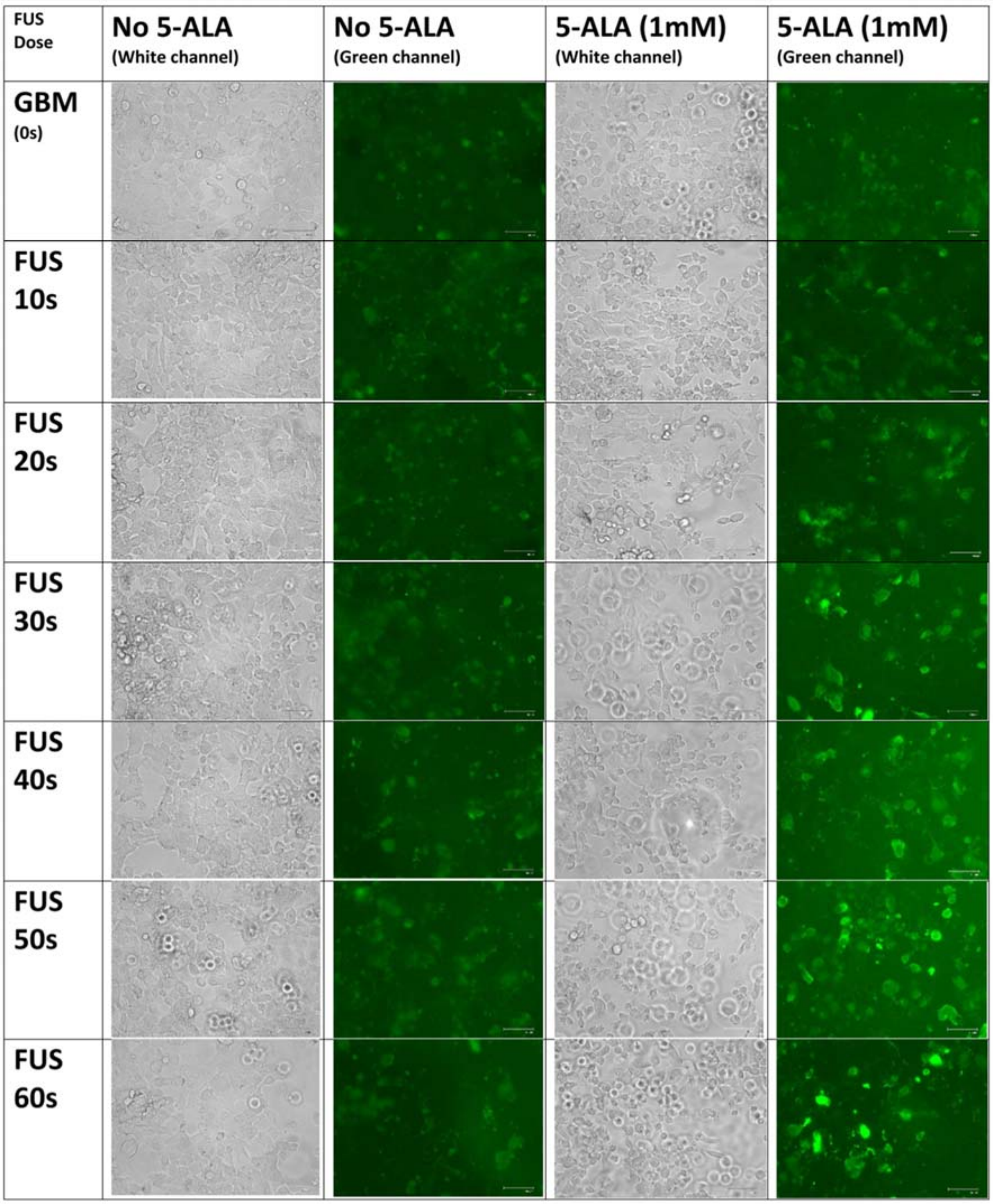


**Figure S7** - Separate green and white channel 40x light microscopy of GBM22 cells treated with Annexin V green, fluorescent apoptosis marker. Conditions no treatment, 5-ALA only (1mM), FUS 10s, 20s, 30s, 40s, 50s cumulative sonication dose (5.5w/cm2, 10% DC, 90ms pulse length), and Sonodynamic therapy at each cumulative sonication dose(5-ALA and FUS). 1.5 hours post-treatment. Scale bar represents 125um. FUS 60s and GBM (0s) panels have been used for Fig 7B.

.

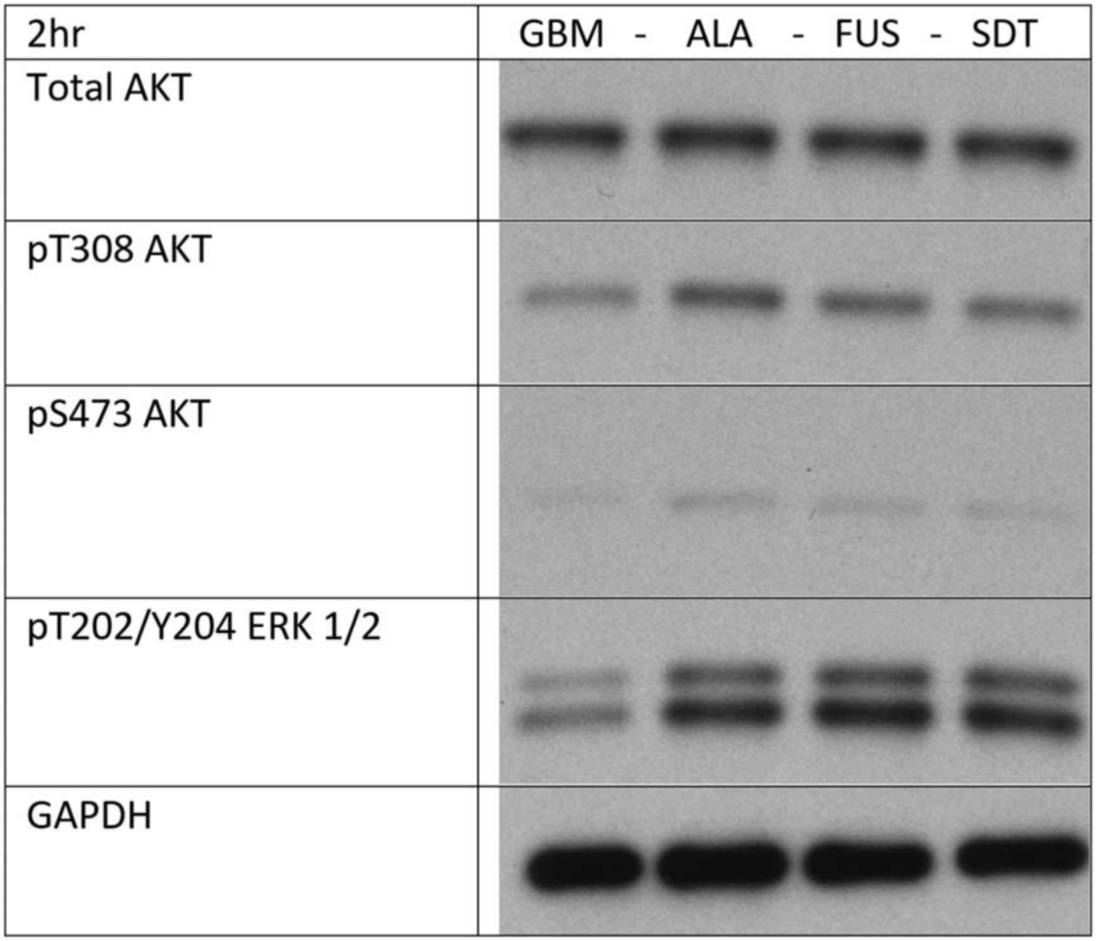


**Figure S8** – Western blot results for AKT, P-AKT (308), P-AKT (473), P-ERK (p40-44), and related GAPDH, lysed 1.5hours post-treatment. Conditions: no treatment, 5-ALA only (1mM), FUS only (0.4w/cm2, 60s cumulative sonication, 10% DC, 90ms pulse length), Sonodynamic therapy (5-ALA and FUS).
